# M1 and M2 macrophages differentially regulate colonic crypt renewal

**DOI:** 10.1101/2022.07.13.499394

**Authors:** Sathuwarman Raveenthiraraj, Griselda Awanis, Marcello Chieppa, Anastasia Sobolewski

## Abstract

The colonic epithelium is the most rapidly renewing tissue in the body and is organized into a single cell layer of invaginations called crypts. Crypt renewal occurs through Lgr5+ gut stem cells situated at the crypt base, which divide, produce daughter cells that proliferate, migrate, differentiate into all the cells required for normal gut function (eg. Goblet cells, enterocytes), and are finally shed into the crypt lumen. In health this rapid renewal helps maintain barrier function next to the hostile gut luminal environment that contains microbes and food. In parallel, the peri-cryptal lamina propria hosts the largest monocyte-derived macrophage population in the human body. Different macrophage phenotypes have been associated with intestinal health/intact barrier function, namely M2 compared to M1 macrophages that indicate inflammation/compromised barrier function. However, the direct effect of different macrophage subtypes have on colonic crypt renewal is not well understood. In this study we have utilized a reductionist 3D *in vitro* co-culture model to determine the regulatory capacity of M1 and M2 macrophages on colonic crypt renewal. We show that colonic crypt proliferation is increased in the presence of M1 or M2 macrophages, while we further demonstrate that a decrease in goblet and tuft cell expression as well as an increase in Lgr5+ stem cell numbers is only achieved through M1-crypt crosstalk in a contact dependent manner.

## Introduction

The intestinal epithelium, lining the innermost layer of the large intestine, plays a crucial role in the physical protection of the underlying tissue from pathogenic threats, commonly encountered in the lumen, where a single-cell thick epithelium is perpetually renewed every 4-5 days (Darwich et al., 2014). Lgr5 (Leucine-rich repeat containing G protein-coupled receptor 5) expressing stem cells at the base of epithelial invaginations, termed crypts, drive epithelial renewal (Barker et al., 2007; Barker et al., 2012). Here, Lgr5+ intestinal stem cells in the colon generate highly proliferative transit amplifying daughter cells which migrate along the crypt-axis and give rise to fully differentiated epithelial cells such as enterocytes, goblet cells, tuft cells and enteroendocrine cells until they are shed into the lumen at the end of their life cycle (Van der Flier and Clevers., 2009; Williams et al., 2015).

To further counteract the looming threat the large microbial presence poses over the colonic epithelium, the underlying lamina propria employs the densest macrophage population in the human body, where blood derived Ly6C+ monocytes are recruited to the submucosa where they then differentiate towards a mature macrophage phenotype (Thursby and Juge., 2017; Bain et al., 2018, Bain et al., 2014). Through their proximal peri-cryptal localization in the lamina propria, these macrophages swiftly apprehend invasive foreign pathogens in a tolerogenic manner, while an escalating inflammatory response is repressed (Smythies et al., 2005; Hine et al., 2019; Gordon et al., 2014; Rescigno and Chieppa., 2005). Macrophages are highly plastic and their phenotypical properties are often influenced through environmental cues within the lamina propria (Viola and Boeckxstans, 2021).

Early studies have broadly defined two distinctive macrophage phenotypes based on their physiology and function commonly known as M1 and M2 macrophages (Mills et al., 2000; Mills et al. 2012, Orecchioni et al., 2019). Here, acute epithelial injury results in the influx of pro-inflammatory and bactericidal subsets of M1 macrophages, while residential macrophages in the steady-state reportedly possess an M2-like macrophage phenotype (Na et al., 2019; Lissner et al, 2015).

Over the last decade, gene signature studies have postulated that the M1 and M2 activation states likely represent the opposite ends of the phenotypical macrophage spectrum (Jablonski et al., 2014; Orrechioni et al., 2019). Here, several studies have demonstrated that M1 macrophages express distinct pro-inflammatory cytokines and chemokines compared to its M2 macrophage counterpart, where the cytokine profile is dominated by the expression of anti-inflammatory associated chemokines such as IL-10 and TGF-β among other (Mantovani et al., 2004; Shapouri-Moghadam et al., 2017; Viola et al., 2019; Rath et al., 2014; Palmieri et al., 2020). Furthermore, it has been established that M1 and M2 macrophages can be defined by their relative expression of CD38 (Jablonski et al., 2015).

The classical role of macrophages in tissue clearance and intestinal immunity has been extensively studied, however, as M1 and M2 macrophages often cohabitate the submucosal space *in vivo*, little is known regarding their respective capacity to engage with the colonic epithelium and their respective contributory role in epithelial renewal (Bain et al., 2018; Smythies et al., 2005; Smith et al., 2011; Martinez and Gordon., 2014).

Indeed, ablation of the macrophage population in the small intestine resulted in the marked reduction of Lgr5+ expressing stem cells and reduced intestinal motility (Sehgal et al., 2018; De Schepper et al., 2018). Furthermore, early work from Pull et al, demonstrated that a subset of activated macrophages are recruited to the site of injury and induce proliferation of epithelial progenitor cells within the crypt, while Skoczek and colleagues further showed that inflammatory monocytes, a macrophage precursor, are recruited and juxtaposed to Lgr5EGFP+ stem cells at the base of colonic crypts upon exposure to *E*.*coli in vivo* and induced an increase in epithelial proliferation *in vitro* (Pull et al., 2005;Skoczek et al., 2014). As cell-to-cell contact between two cell types may evoke a signaling cascade in the target cell, collectively, these studies suggest that macrophages likely function as a secondary mediator of the intestinal stem cell niche, while it further begs the question as to whether the phenotypic states of M1 and M2 macrophages commonly exhibited during intestinal inflammation and steady-state respectively, can differentially regulate colonic crypt renewal.

The intestinal lamina propria plays an essential role in the maintenance of the colonic stem cell niche, where the underlying mesenchymal, immune cells or extracellular matrix compartments were demonstrated to modulate the stem cell niche (McCarthy et al., 2020; Shoshkes-Carmel et al.,2018; Meran et al., 2017; Onfroy-Roy et al., 2020). However, due to the myriad of subepithelial signaling stimuli involved, *in vivo* models face challenges in delineating their respective effects on the stem cell niche. As most adult intestinal macrophages are derived from the monocytic cell lineage, we are able to mirror the *in vivo* crypt-macrophage microenvironment using our *in vitro* reductionist 3D co-culture model, allowing for the close spatial-temporal study of bone-marrow derived M1 and M2 macrophage interactions and its effects on colonic crypt renewal (Skozcek et al., 2014; Bain et al., 2014).

We show that both M1 and M2 macrophage can increase colonic crypt proliferation, while M1 macrophages further induce colonic crypt proliferation through secretory factors. We further demonstrate that juxtracrine contact between M1, but not M2 macrophages results in decreased tuft and goblet cell expression whilst observing an increase in Lgr5 expressing stem cell numbers, where direct M1 macrophage-epithelial interactions result in the upregulation in downstream Wnt-signaling targets LEF1 and CyclinD1 in the colonic epithelium.

## Results

### M1 and M2 macrophages stimulate the proliferation of colonic crypts

To determine the effects M1 and M2 macrophages have on colonic crypt growth, we cultured M1 or M2 macrophages along with freshly isolated crypts, where macrophages are either in contact or proximally localized near the crypts **(Fig 1A)**. Primarily macrophages were found to be in contact with the base and mid region of the crypt **(Supplementary Fig 2)**. We then examined EdU incorporation in the colonic epithelium using immunofluorescent microscopy (**Fig. 1B**). The co-culture of crypts with either M1 or M2 macrophages resulted in a significant increase of EdU incorporation (green) compared to control (**Fig 1B**). Most notably, EdU incorporation was also significantly higher in crypts cultured with M1 macrophages when compared to M2 macrophages. Further analysis of epithelial caspase-3 expression (**Supplementary Fig. 1**) did not show any significant changes between crypts cultured with M1 or M2 macrophages compared to control, while morphological analysis of colonic crypt length signified a shortening of crypt length in crypts cultured with M1 macrophages compared to control crypts (**Supplementary Fig. 2C)**.

**Figure 1:**
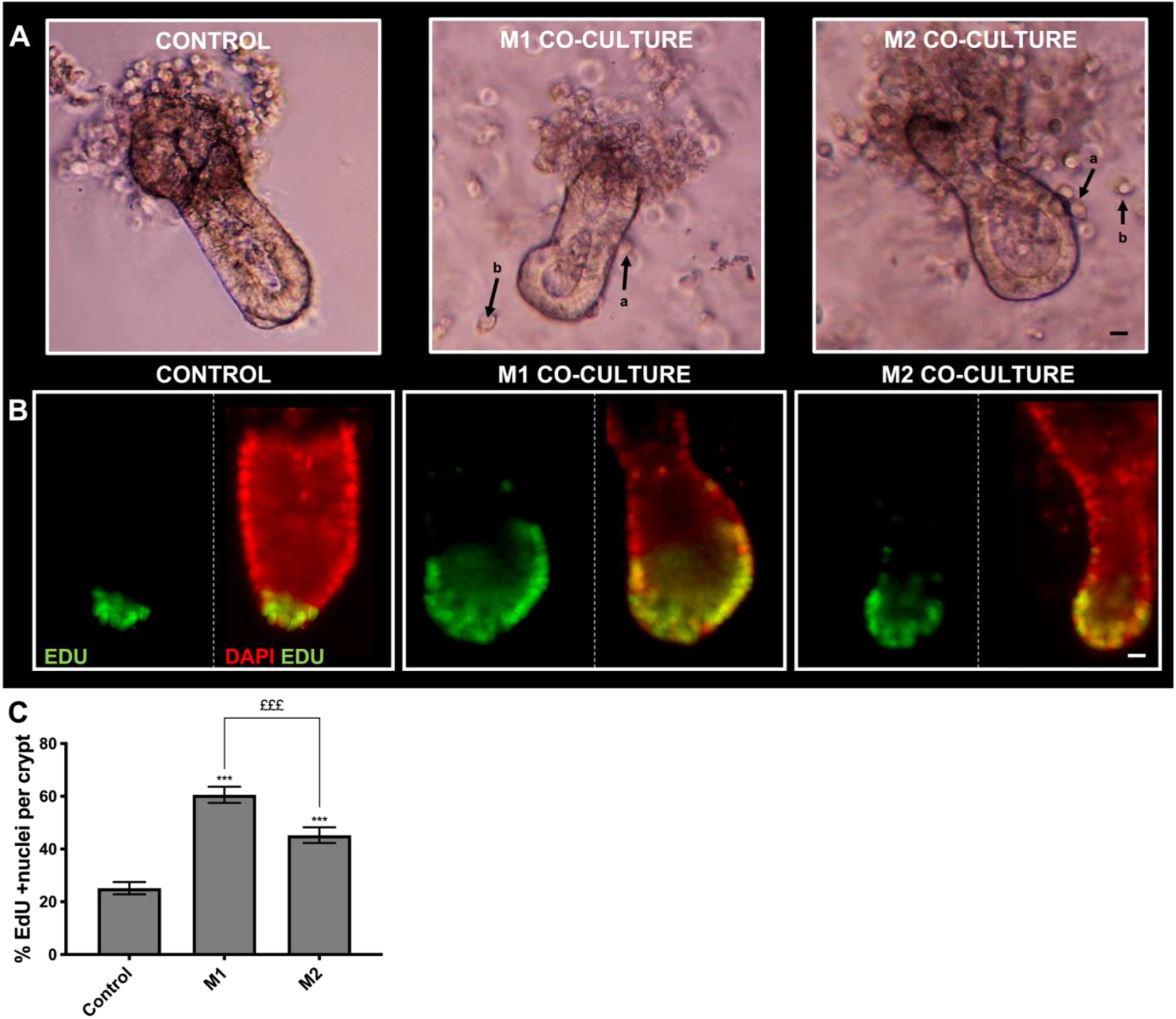
M1 and M2 macrophages increase EdU incorporation of colonic crypts in in vitro co-culture. **A)** Representative white light images showing crypts cultured alone and with M1 or M2 macrophages, where macrophages are either **a)** in contact or **b)** not in contact with crypts (white arrows). Scale bar at 15µm **B)** Representative epi-fluorescent images showing EdU incorporation (green) in the nuclei (red) within colonic crypt-macrophage co-cultures. Co-labelling of nucleus and EdU shown in yellow **C)** Histogram showing the percentage of EdU positive nuclei per crypt within the macrophage subtype co-culture condition. (n=3, ***P< 0.001 compared to control; M1 compared to M2 £££ P< 0.001). Scale bar at 15µm.

### M1 macrophages, but not M2 macrophages reduce goblet and tuft cell numbers in colonic crypts

To determine whether M1 or M2 macrophage affect the differentiated cell compartment within colonic crypts, we aimed to quantify enteroendocrine, tuft and goblet cell numbers using confocal microscopy (**Fig 2**). Using chromogranin-A (white) and E-cadherin (red), we visualized Cro-A + enteroendocrine cells present in crypts cultured with M1 or M2 macrophages compared to control (**Fig 2A**). Here we show that the culture of M1 or M2 macrophage with crypts does not significantly affect enteroendocrine cell numbers (**Fig 2B**).

**Figure 2:**
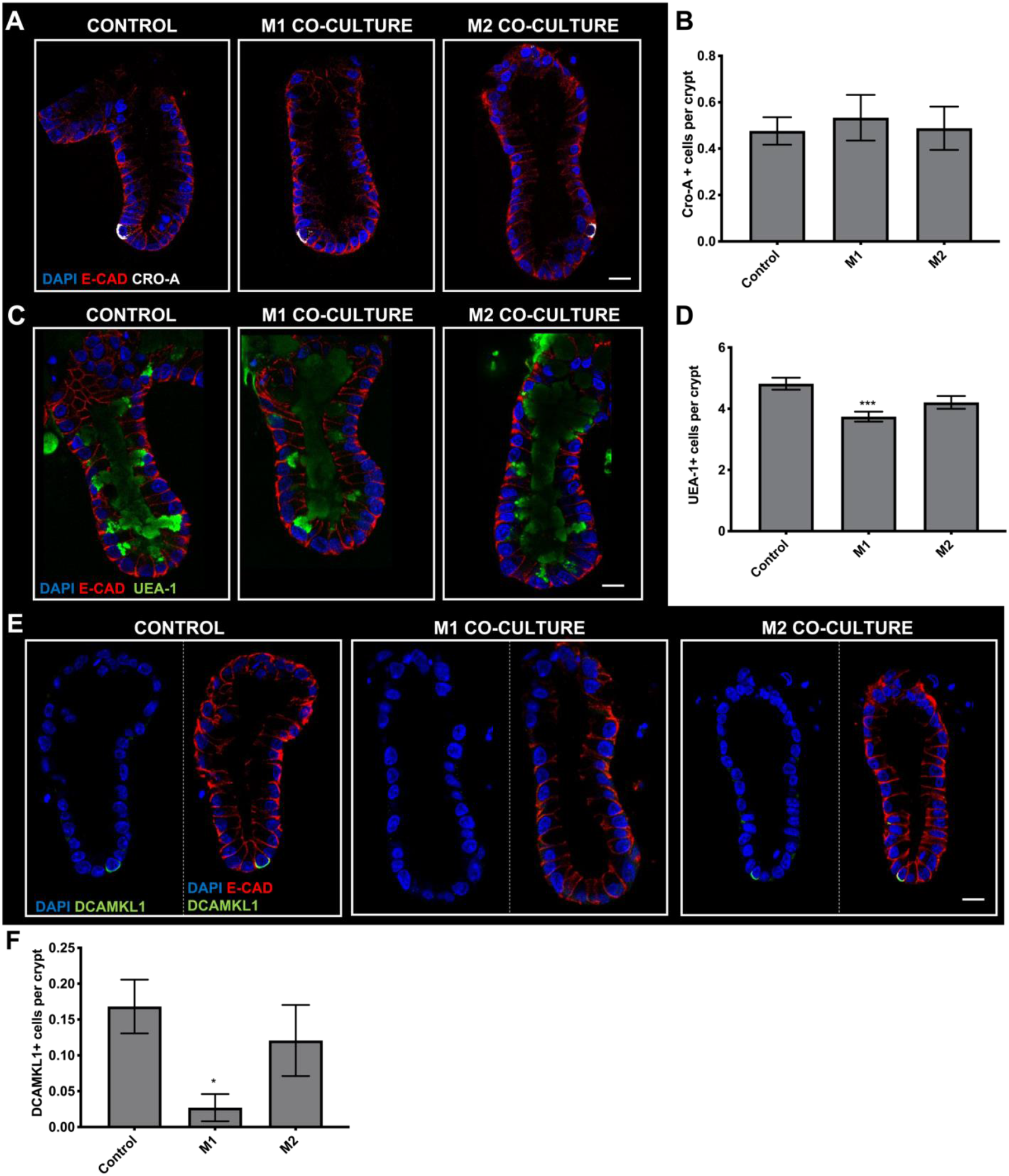
Cro-A+ cell numbers are maintained in crypts cultured with M1 or M2 macrophages, while UEA-1+ goblet cells and DCAMKL1+ tuft cell numbers are decreased in crypts cultured with M1 macrophages but not M2. **A)** Representative confocal images showing Chromogranin-A(Cro-A) expression (white), DAPI (blue) and E-cadherin (red) in crypt-macrophage subtype co-culture. **B)** Histogram showing the average number of Cro-A positive cells per crypt within each co-culture condition (n=4, ns). **C)** Representative confocal images showing UEA-1 expression (green), DAPI (blue) and E-cadherin (red) in each crypt-macrophage subtype co-culture **D)** Histogram showing the average number of UEA-1 positive cells per crypt within each co-culture condition (n=5, **P <0.01 ***P<0.0001 compared to Control). **E)** Representative confocal images showing DCAMKl1 expression (green), DAPI (blue) and E-cadherin (red) in each crypt-macrophage subtype co-culture **F)** Histogram showing the average number of DCAMKL1 positive cells per crypt within each co-culture condition (n=5, *P<0.05 compared to Control). Scale bar at 20μm.

Next, we used the *Ulex europaeus agglutinin-* UEA-1 (green) and E-cadherin (red) to identify goblet cells within crypts cultured with M1 and M2 macrophages compared to control (**Fig 2C**). We show that the co-culture of M1 macrophages with colonic crypts results in a significant decrease in UEA-1+ goblet cell numbers compared to control, while crypts cultured with M2 macrophage maintained UEA-1+ goblet cell numbers (**Fig 2D**).

To determine whether the tuft cell numbers were affected in crypts cultured with M1 or M2 macrophages, we visualized the epithelial tuft cell population using DCAMKL1 (green) and E-cadherin (red) (**Fig 2E**). Here, we observed a significant reduction in DCAMKL1+ tuft cell numbers in crypts cultured with M1 macrophages compared to control, while DCAMKL1+ tuft cells were maintained in crypts cultured with M2 macrophages (**Fig 2F**).

### M1 macrophages increase Lgr5+ stem cell numbers in colonic crypts

Having shown the effect of M1 and M2 macrophages on the differentiated cell population, we wanted to further understand their effect on the epithelial stem cell population. Here we identified the colonic stem cell population using the leucine-rich G-protein coupled receptor 5, Lgr5 (green) and E-cadherin (red) (**Fig 3A**). We observed a significant increase in Lgr5+ stem cell numbers in crypts cultured with M1 macrophages compared to control, while Lgr5+ stem cell numbers were comparable to control in crypts cultured with M2 macrophages (**Fig 3B**). Further analysis of Lgr5+ stem cell position within the colonic crypt compartment has shown that an increased number of Lgr5+ stem cells were localized in the base and mid region of crypts cultured with M1 macrophages when compared to control and crypts co-cultured with M2 macrophages. No significant changes were noted in the top region of colonic crypts cultured with either M1 or M2 macrophages compared to control (**Fig 3C**).

**Figure 3:**
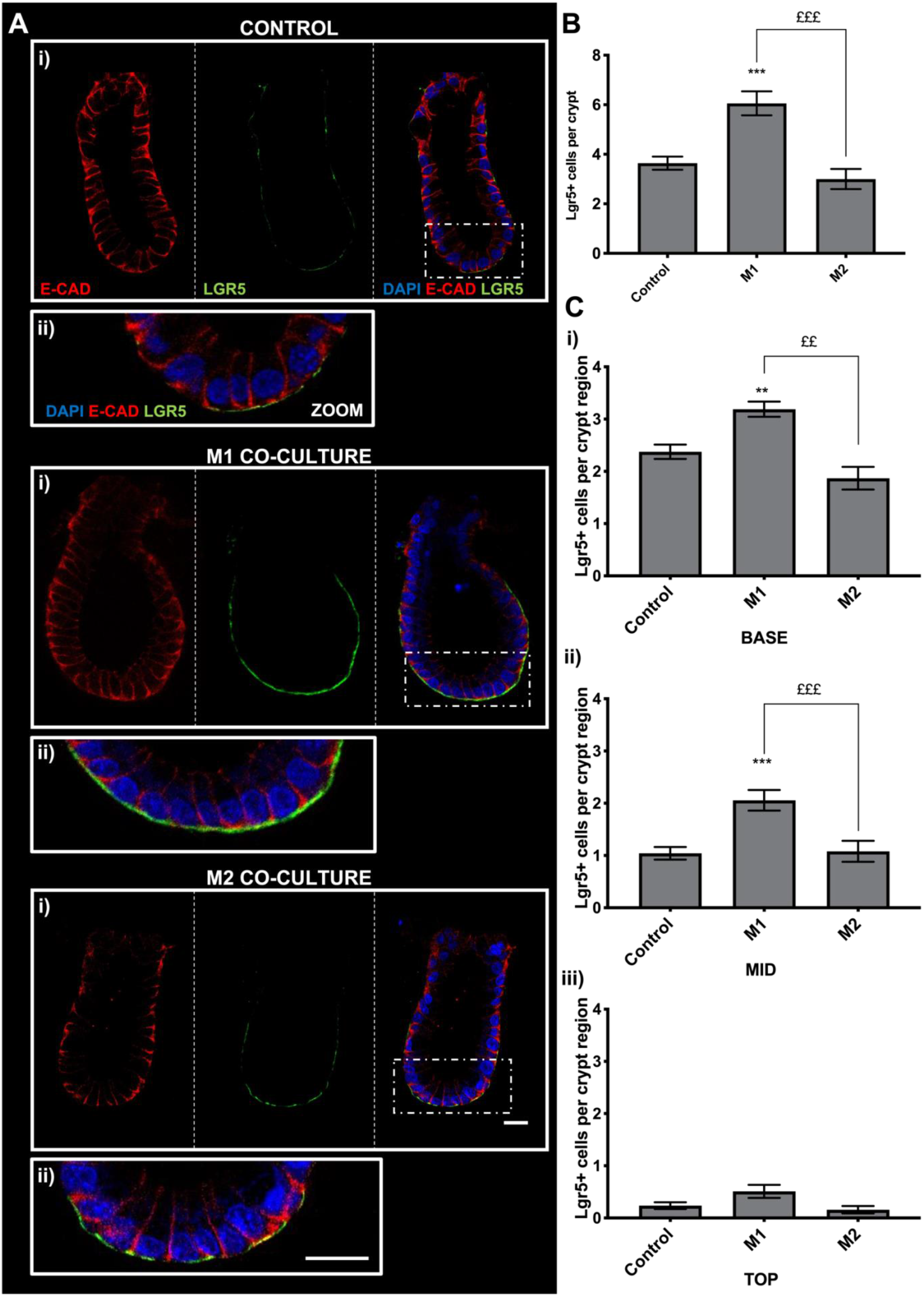
M1 but not M2 macrophages increase in vitro Lgr5+ cell expression in colonic crypts within the co-culture model. **A) i)** Representative confocal images showing Lgr5 expression (green), DAPI (blue), E-cadherin (red) and brightfield (white) in each crypt-macrophage subtype co-culture and **ii)** enlarged image of crypt base **B)** Histogram showing the average number of LGR5 positive cells per crypt within each co-culture condition **C)** Histogram showing the position of Lgr5 positive cells within each crypt region **i)** base, **ii)** mid and **iii)** top). (n=4, **P<0.01 compared to control, ***P<0.001; M2 compared to M1 £££ P<0.001; ££ P< 0.01). Scale bar 20μm.

### Epithelial proliferation is increased via secretory factors in M1 macrophages while M2 macrophages require juxtracrine-contact

As we have shown that EdU incorporation significantly increased in crypts cultured with M1 and M2 macrophages, we next determined whether the previously observed effects stem from physical cell-cell contact between macrophages and colonic crypt cells as observed *in vitro* (**Supplementary Fig. 2B**) or co-culture derived secretory products using a conditioned media model (**Supplementary Fig 3A**). Here we show representative images of EdU (green) incorporation in crypts cells (red) cultured in the presence of M1 or M2 macrophages (co-culture) and crypts cultured without direct contact to M1 or M2 macrophages (CM) but placed in vicinity to a co-culture (**Fig 4A**). Semi-quantitative analysis of the percentage of EdU+ positive cells compared to the total number of DAPI positive cells shows that crypts directly cultured with M1 and M2 macrophages a significantly increases EdU incorporation when compared to control crypts (**Fig 4B**). When crypts were cultured without direct contact with M1 macrophages (M1-CM) but placed in vicinity to an M1-crypt co-culture, the percentage of EdU incorporation was significantly increased compared to control crypts. No significant changes in the percentage of EdU incorporation were found in crypts cultured without direct contact with M2 macrophages (M2-CM) but placed in vicinity to the M2-crypt co-culture compared to control crypts.

**Figure 4:**
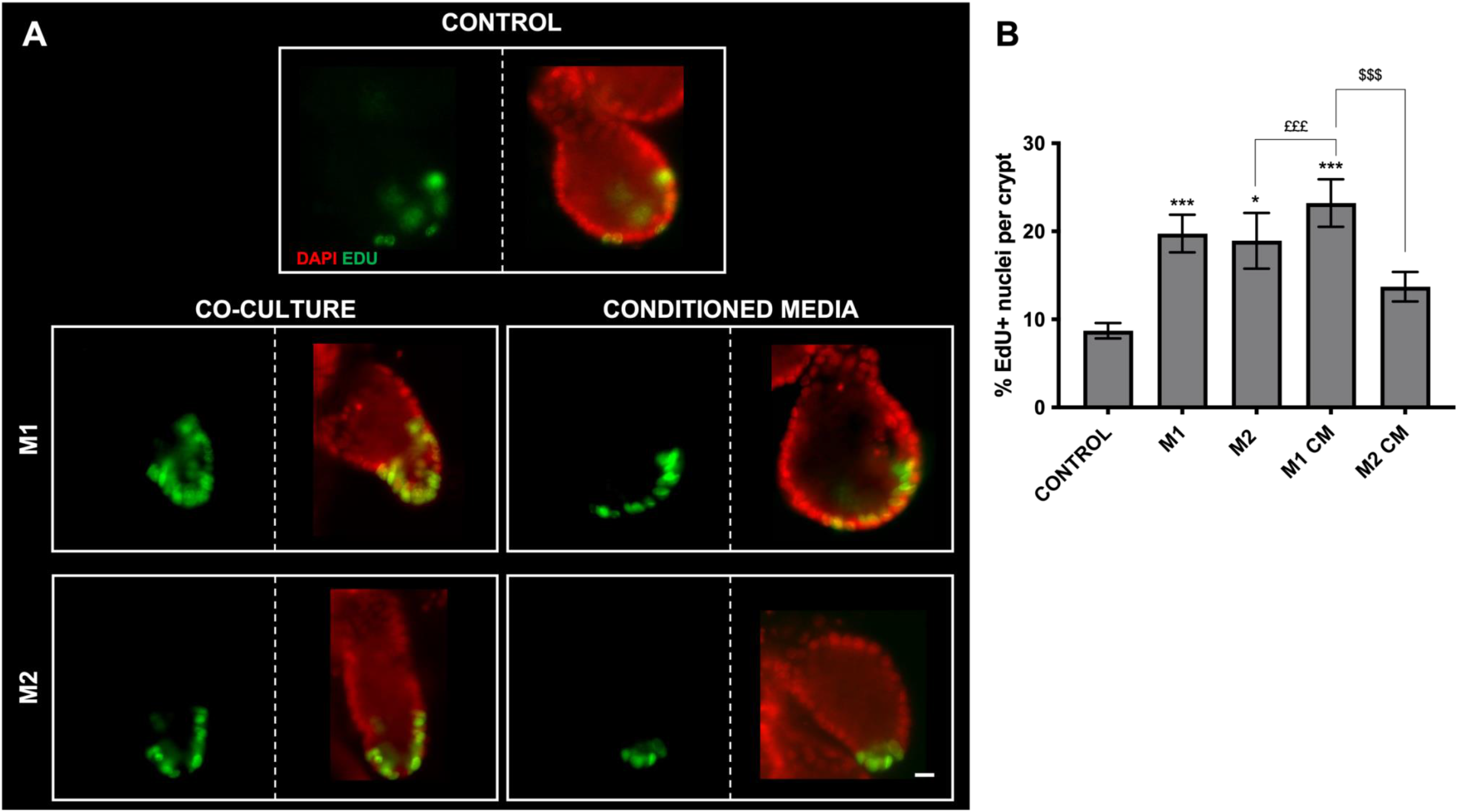
Juxtracrine contact is utilised by both M1 and M2 macrophages to induce increased EdU incorporation in colonic crypts, while M1 macrophages can further increase EdU incorporation via secretory factors in vitro. **A)** Representative epi-fluorescent images showing EdU incorporation (green) in the nuclei (red) within colonic crypt-macrophage co-culture and conditioned media (CM). **B)** Histogram showing the percentage of EdU positive nuclei per crypt within each co-culture and conditioned media (CM) culture model. (n=4, compared to Control *P<0.05, **P<0.01, ***P<0.001; M1 compared to M2 $$P<0.01) Scale bar at 15μm.

### M1 macrophage-epithelial juxtracrine contact is required to reduce goblet and tuft cell numbers

To determine whether the reduction in UEA-1+ goblet cell and DCAMKL1+ tuft cell numbers in crypts cultured with M1 macrophages were induced via physical contact or co-culture derived secretory factors we devised a conditioned media model (**Supplementary Fig 3B**). We show representative confocal images of crypts cultured alone, cultured along with M1 macrophages (M1 co-culture), crypts cultured without direct contact to an M1-crypt co-culture (M1-CM) and crypts cultured alone but placed in vicinity to M1 macrophages (M1-only) (**Fig 5A**). UEA-1+ goblet cells were identified and analyzed using UEA-1 (green) and E-cadherin (red) where we show that their numbers in crypts cultured under M1 CM and M1-only culture models were maintained. However, a significant decrease in the number of UEA-1 positive cell expression was observed in crypts directly cultured in the presence of M1 macrophages compared control crypts (**Fig 5B**). A significant reduction in UEA-1+ goblet cells numbers was noted in the mid region of the crypts when cultured with M1 macrophages compared to control crypts.

**Figure 5:**
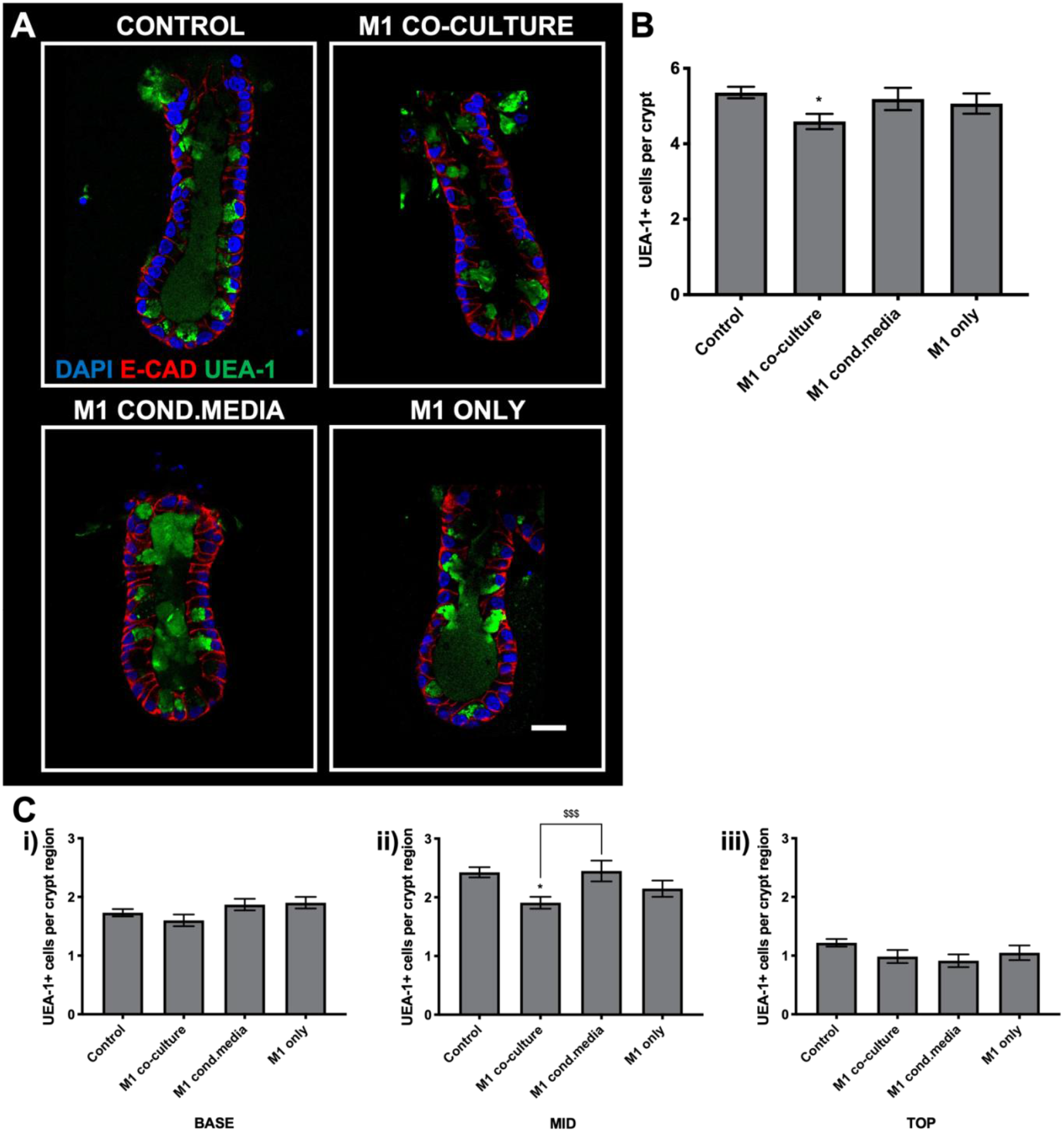
Physical contact between M1 macrophages and colonic crypts but not secretory factors decrease DCAMKL1+ cell expression in crypts in vitro. **A)** Representative confocal images showing UEA-1 expression (green), nuclei (blue) and E-cadherin (red) in crypts cultured in the M1 co-culture, M1 -CM media and M1-only culture models. **B)** Histogram showing the average number of UEA-1 positive cells per crypt cultured in the M1 co-culture, M1-CM and M1-only culture models **C)** Histogram showing the average number of UEA-1 positive cells per crypt region **i)** base, **ii)** mid and **iii)** top when cultured in the M1 co-culture, M1-CM and M1 only culture models. (n=6, *P<0.05 compared to Control; $$$P<0.001 compared to M1 co-culture) Scale bar 20μm.

In a similar manner, we determined whether direct cell to cell contact between macrophages and colonic crypts is required to induce a reduction in DCAMKL1+ tuft cell number. Here we show a significant reduction in DCAMKL1+ (green) tuft cell numbers in crypts cultured with M1 macrophages (**Fig 6A**). In comparison, the number of DCAMKL1+ cells in crypts of M1-CM and M1-only culture model were maintained compared to control crypts (**Fig 6B**). Furthermore, a significant reduction in DCAMKL1+ tuft cell number was observed at the base region of the colonic crypt when cultured with M1 macrophages compared to control crypts.

**Figure 6:**
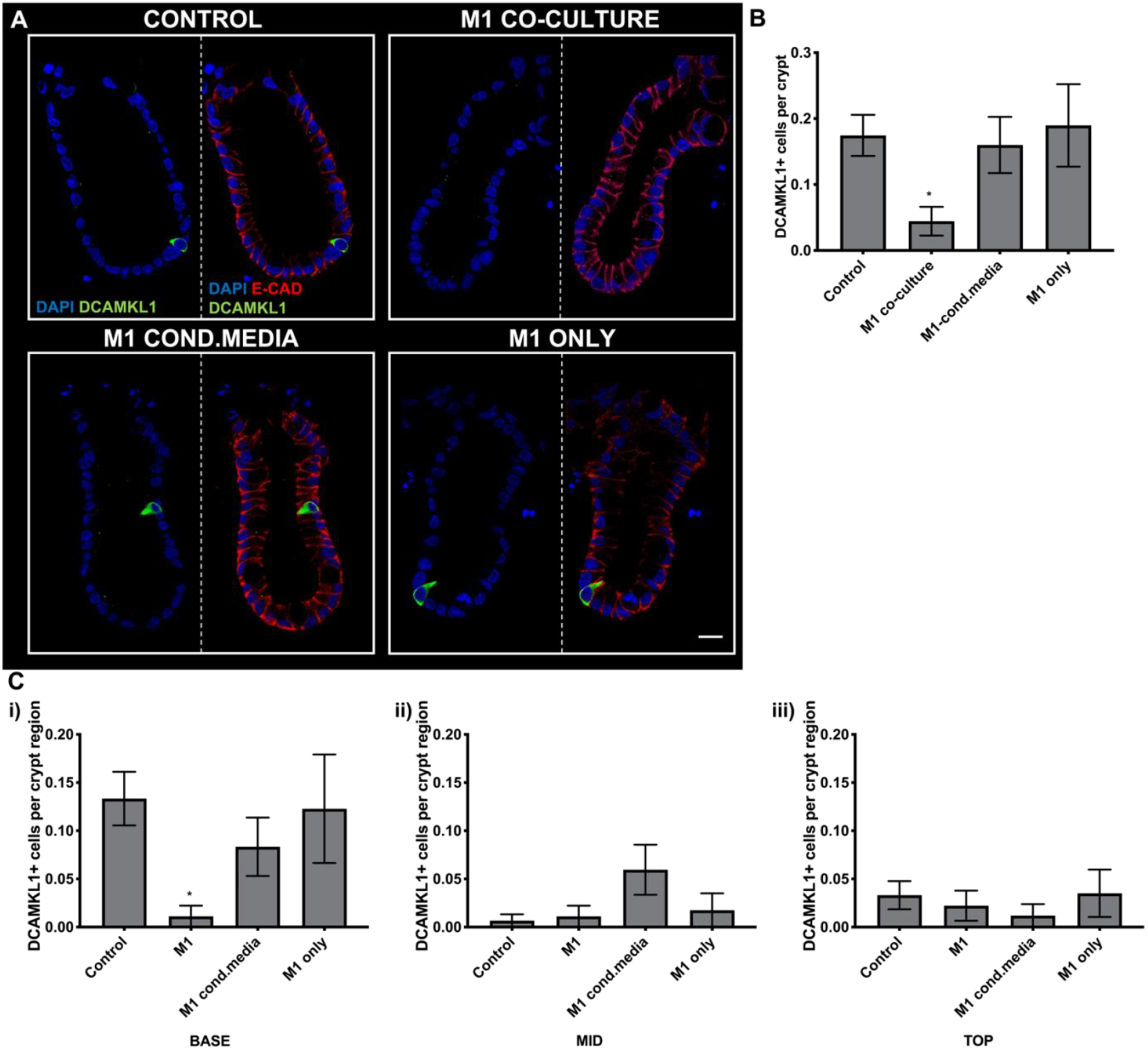
Figure 6.3: Physical contact between M1 macrophages and colonic crypts but not secretory factors decrease DCAMKL1+ cell expression in crypts in vitro. **A)** Representative confocal images showing DCAMKL1 expression (green), nuclei (blue) in crypts cultured in the M1 co-culture, M1-CM and M1-only culture models. **B)** Histogram showing the average number of DCAMKL1 positive cells per crypt cultured in the M1 co-culture, M1-CM and M1-only culture models **C)** Histogram showing the position of DCAMKL1+ cells per crypt region **i)** base **ii)** mid **iii)** top in M1 co-culture, M1-CM and M1-only culture models. (n=6, *P<0.05 compared to Control) Scale bar 20μm.

### Juxtracrine contact between M1 macrophages and colonic crypts is required to increase Lgr5+ stem cell expansion

We have previously established that M1 macrophages increase Lgr5+ stem cell expression in colonic crypts. To understand the mode of mechanism required to bring about such changes, we next determined whether proximal macrophage-crypt contact, or secretory factors are required to increase crypt stem cell expression. We show representative confocal images of crypts cultured alone, cultured along with M1 macrophages (M1 co-culture), crypts cultured without direct contact to a M1-crypt co-culture (M1-CM) and crypts cultured alone but placed in vicinity to M1 macrophages (M1-only), where we identified the stem cell population using Lgr5 (green) and E-cadherin (red) (**Fig 7A**). The number of Lgr5 positive cells per crypt was significantly increased in crypts cultured in the presence of M1 macrophages compared to control crypts. However, in crypts of M1-CM and M1-only culture models, Lgr5 positive cell numbers were maintained compared to control. Furthermore, crypts cultured along with M1 macrophages also expressed significantly more Lgr5 positive cells in comparison to crypts within the M1-CM and M1-only culture models (**Fig 7B**).

**Figure 7:**
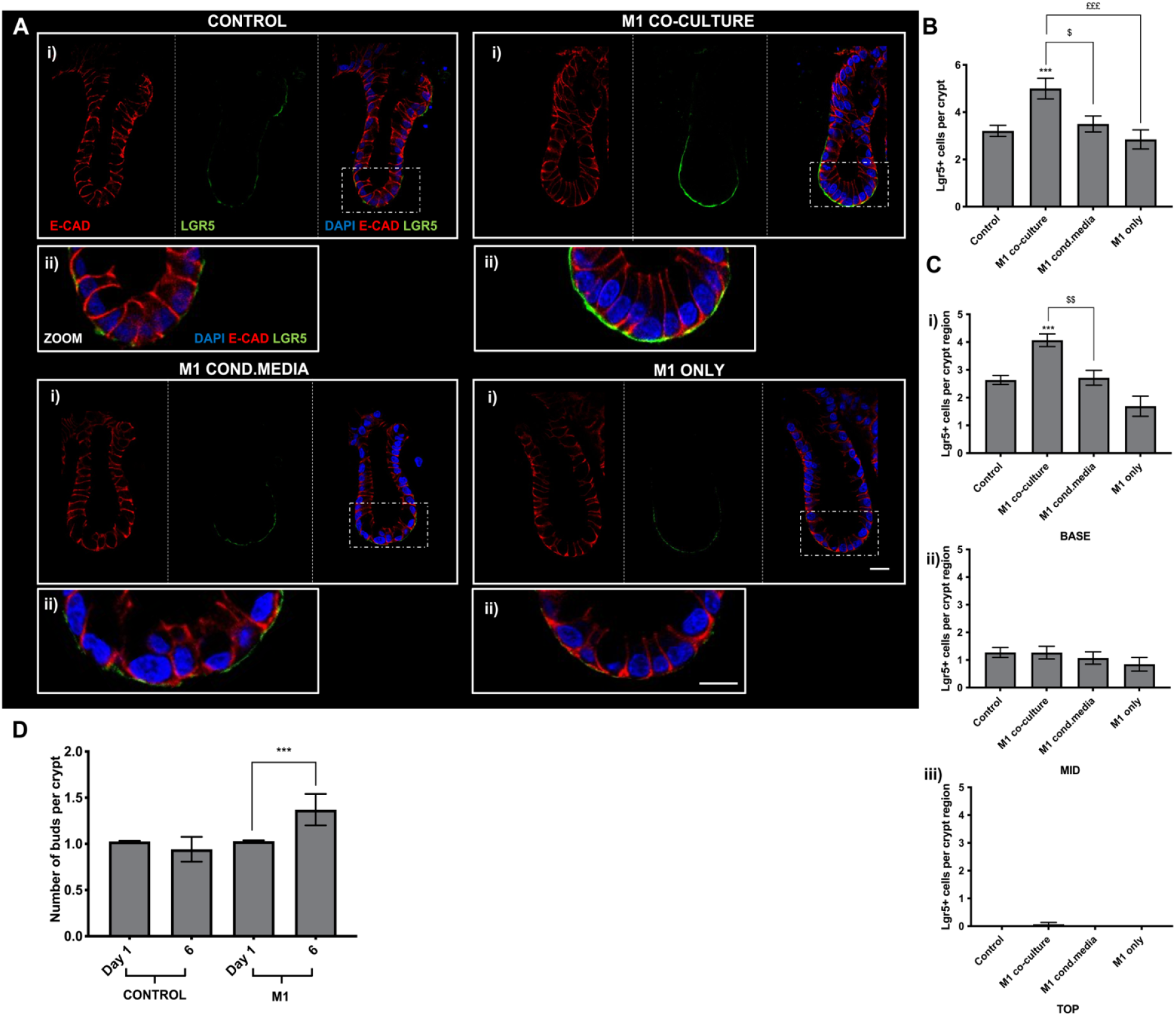
M1 macrophages induce an increase in Lgr5+ cell expression via physical contact; not secretory factors in colonic crypts in vitro. **Ai)** Representative confocal images showing basal Lgr5 expression (green), nuclei (blue) and E-cadherin (red) in crypts from M1 co-cultures, M1-CM and M1-only culture models. **Aii)** Enlarged confocal images showing expression of Lgr5 (green) along the base of the crypt alongside white light or DAPI (blue), E-cadherin (red) when cultured in M1 co-culture, M1-CM or M1-only models. **B)** Histogram showing the average number of Lgr5 positive cells per crypt cultured in M1 co-culture, M1 and M1 only culture models. **C**) Histogram showing the position of Lgr5+ cells per crypt region **i)** base, **ii)** mid and **iii)** top when cultured in M1 co-culture, M1-CM and M1-only models. (n=3, ***P<0.001 compared to Control; $P<0.05, $$P<0.01 M1 co-culture compared to M1-CM; £££P<0.001 M1 co-culture compared to M1-only). **D)** Histogram showing the number of buds per crypt expressed on Day 1 and 6 days in control crypts compared to M1-crypt co-culture (n=3; Control vs M1 ***P<0.0001). Scale bar at 20µm.

Further analysis of the distribution of Lgr5 throughout each crypt compartment has shown that the majority of Lgr5 positive cells were located at the base of the crypts with fewer located at the mid region and the lowest numbers found at the top. Significantly more Lgr5 positive cells were expressed at the base of the crypt cultured with M1 macrophages, where numbers in the mid and top region were similar compared to control crypts. The number of Lgr5 positive cells expressed at the base, mid and top region of crypts in the M1-CM and M1-only model were also comparable to control crypts (**Fig 7C**). Furthermore, long term culture of M1 macrophages with colonic crypts resulted in increased colonic crypt budding following 6 days in culture compared to control crypts (**Fig 7D**).

### Juxtracrine-contact is required to increase downstream Wnt targets proteins LEF1 and CyclinD1

To determine whether physical cell-cell contact between macrophages and colonic crypts, or co-culture derived secretory factors differentially affect the Wnt target proteins, expression of CyclinD1 and LEF1 was further studied. LEF1 expression (red) is expressed within DAPI (blue) positive cells in colonic epithelial cells in control crypts, crypts cultured without direct contact to M1 macrophages but placed in vicinity to a M1-crypt co-culture (M1-CM) and crypts cultured without direct contact to M1 macrophages but placed in vicinity of M1 macrophages alone (M1-only) (**Fig. 8A**).Semi-quantitative analysis of the mean fluorescence intensity of LEF1 expression per crypt region is shows the mean fluorescence intensity of LEF1 expression in control crypts is shown to be evenly distributed along the longitudinal crypt-axis, in comparison, LEF1 expression in crypts cultured in the presence of M1 macrophages was significantly higher at the base, mid and top region of the crypts compared to control, M1-CM and M1-only crypts (**Fig 8B**).

**Figure 8:**
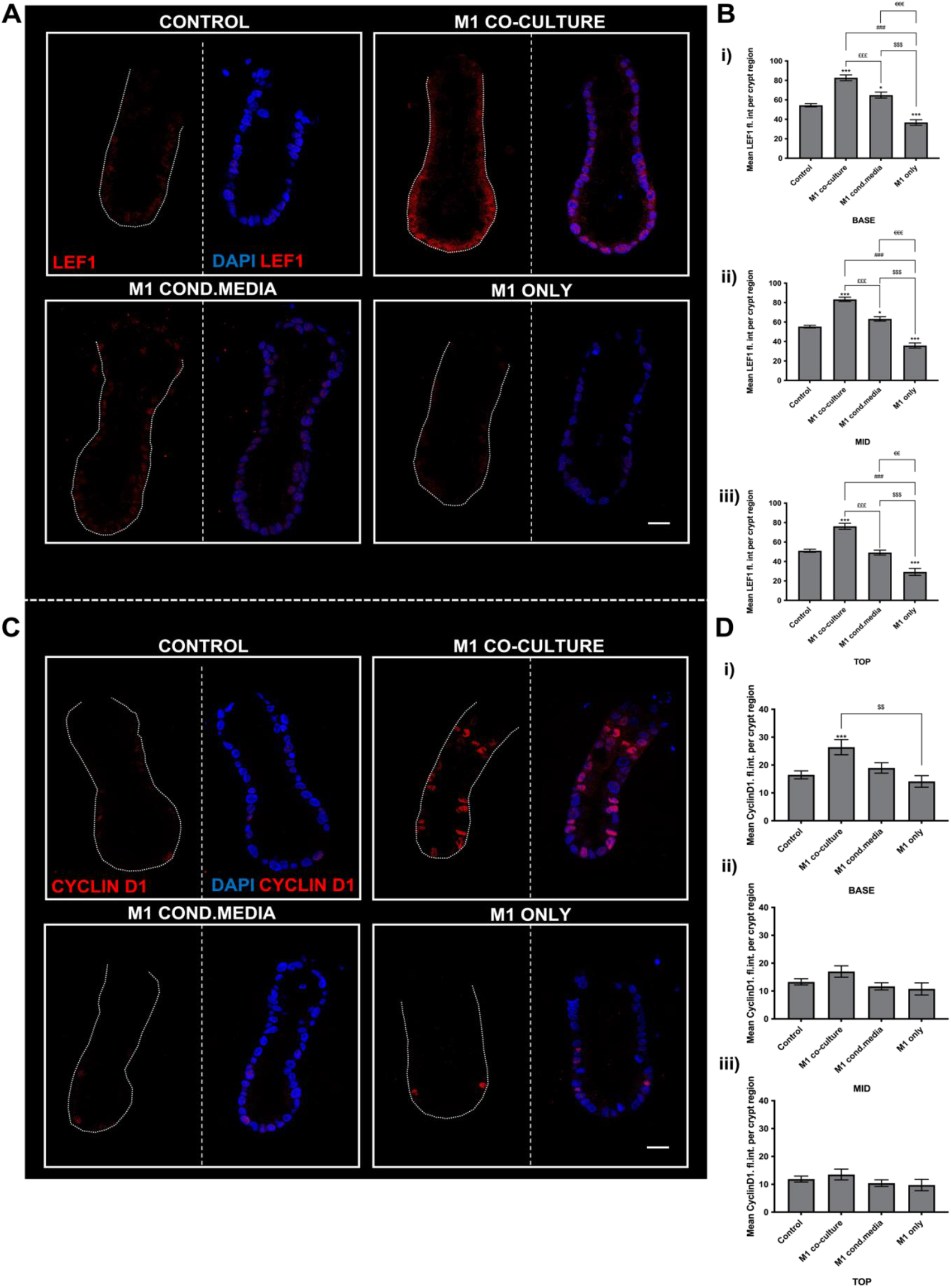
M1 macrophages induce an increase in colonic LEF1 expression through physical contact and secretory factors, while colonic CyclinD1 expression is increased through physical contact but not secretory factors. ***A)*** *Representative confocal images showing nuclear LEF1 expression (red), nuclei (blue) in crypts from M1 co-cultures, M1-CM and M1-only culture models* ***B)*** *Histogram showing the average fluorescence intensity of LEF1 within each crypt co-culture/conditioned media experiment per crypt region* ***i)*** *base* ***ii)*** *mid* ***iii)*** *top (n=3, ***P<0*.*001 compared to Control; £££P<0*.*001 M1 co-cultured compared to M1-CM; $P<0*.*05,$$P<0*.*01,$$$P<0*.*001 M1 co-culture compared to M1 only. Scale bar at 20µm*. ***C)*** *Representative confocal images showing nuclear Cyclin D1 expression (red), nuclei (blue) in crypts from M1 co-cultures, M1-CM and M1-only culture models*. ***D)*** *Histogram showing the average fluorescence intensity of Cyclin D1 within each crypt co-culture/conditioned media experiment per crypt region* ***i)*** *base* ***ii)*** *mid* ***iii)*** *top. (n=3, compared to Control*. (n=3, ***P<0.001 compared to Control; $$P<0.01 M1 co-culture compared to M1 only. Scale bar at 20 μm.

Similarly, the expression of Cyclin D1, in each co-culture condition was determined (**Fig. 8C**). In control crypts, similar expression of CyclinD1 was found at the base, mid and top of the crypt. In crypts cultured in the presence of M1 macrophages (M1 co-culture) a significant increase in CyclinD1 expression was noted at the base of the crypt compared to control crypts, however CyclinD1 expression was maintained at the mid and top region of crypts. In crypts cultured without direct contact to M1 macrophages but placed in vicinity of a M1-crypt co-culture (M1 CM), the mean fluorescence intensity of CyclinD1 expression remained unchanged at the base, mid and top region of the crypt compared to control crypts. In crypts cultured without direct contact to M1 macrophages but placed in vicinity of M1 macrophages alone (M1-only), the mean fluorescence intensity of CyclinD1 expression also remained unchanged compared to control crypts. Notably, the mean CyclinD1 fluorescence intensity in crypts cultured in the presence of macrophages (M1 co-culture) were significantly higher compared to crypts from the M1-only culture model (**Fig 8D**).

## Discussion

Our study demonstrates that M1 and M2 bone-marrow differentiated macrophages differentially regulate colonic crypt renewal. Epithelial proliferation is increased by either subset with M2 macrophages requiring crypt cell contact and M1 inducing growth through a secreted factor. M2 macrophages maintain intestinal stem cell and differentiated cell numbers throughout the colonic crypt, while M1 macrophages reserve the unique ability to trigger an increase in Lgr5+ stem cells, as well as reduce UEA-1+ goblet cell and DCAMKL1+ tuft cell numbers in a juxtracrine-contact dependent manner. Furthermore, these M1-induced changes are accompanied by the upregulation of downstream Wnt-signaling targets LEF1 and CyclinD1.

While previous studies have shown that macrophages can upregulate intestinal epithelial proliferation in murine *in vivo* injury models, we are first to demonstrate that macrophages can directly increase epithelial proliferation in healthy colonic crypts *in vitro* (Sehgal et al., 2018; Quiros et al., 2017; Pull et al., 2005; Morhardt et al., 2019). Indeed, M1 macrophage-secretory factors such as IL-6, iNOS and HIF-1α previously been identified as known triggers of epithelial proliferation (Rath et al., 2014; Kuhn et al., 2014; Jeffery et al., 2016; Kato et al., 2016; Goggins et al., 2021). As contrasting metabolic signatures and cytokine profile has been attributed to the M1 and M2 macrophage phenotype, we therefore postulate that these contribute to the differential epithelial proliferation we have observed in our study and will be subject of future work (Viola et al., 2019).

We demonstrate that juxtracrine interactions of the epithelium with M1 macrophages results in the reduction of goblet cells numbers, whereas M2 macrophages can maintain goblet cell numbers within the crypt-macrophage co-culture model. UEA-1+ mucus-producing goblet cells play a vital role in intestinal homeostasis, where ablation of goblet cells resulted in spontaneous colitis in mice, while goblet cell expression and its mucosal products were severely altered in patients with IBD (Parikh et al., 2019; Johansson et al., 2014; Lissner et al., 2015). Interestingly, it has also been demonstrated that the ablation of macrophages in the small intestine, results in increased goblet cell numbers, suggesting that goblet cell differentiation fate can be regulated by intestinal macrophages. Vice versa other studies have shown a reduction in goblet cell expression in inflammatory bowel diseases, where the M1 macrophage phenotype is ubiquitously represented (Sehgal et al., 2018; Lissner et al., 2015; Gersemann et al., 2009; Strugala et al., 2008).

M1 macrophages also decreased the number of DCAMKL1+ tuft cells, in colonic crypts through juxtracrine-contact, while crypt tuft cell numbers were maintained in the presence of M2 macrophages. The role of tuft cells within the intestinal epithelium is yet to be fully understood, however colonic *in vivo* studies have shown that ablation of *Atoh1* a downstream Notch signaling transcription factor, resulted in the depletion of DCMAKL1+ tuft cells in the colon, possibly suggesting that M1 macrophages may utilize the Notch signaling pathway to suppress tuft cell differentiation (Herring et al., 2018). Interestingly, a decrease in tuft cell numbers was reported in patients with ulcerative colitis, an intestinal disease in which the activated M1 macrophages play an active role in disease progression (Kjærgaard et la., 2021; Dharmasiri et al., 2021). However, it is unclear whether macrophages can directly inhibit tuft cell expression or whether alterations in the Notch signaling cascade resulted in the depletion of tuft cell differentiation within the colonic crypt. Therefore, further work must be undertaken to delineate the functional significance of tuft cells within the intestinal epithelium, which remains challenging due to the rare occurrence of this epithelial cell type (McKinley et al., 2017).

Within the secretory cell lineage, Cro-A+ enteroendocrine cell numbers were maintained throughout the epithelium in our study. The differentiation of enteroendocrine cells from crypt progenitor cells requires the expression of *Atoh1* and *Neurogenin3* and as studies have also shown that enteroendocrine cells are a highly conserved population within the intestinal epithelium, our findings suggests that it is unlikely that macrophages are able to influence the enteroendocrine cell fate (Hartenstein et al., 2011; Reuters et al., 2022) Instead, it is likely that enteroendocrine cell-derived hormones and peptides may alter macrophage function and further studies should investigate the potential impact these crosstalk mechanism may have in gastrointestinal tract disorders (Worthington et al., 2018).

Previous work from Sehgal and colleagues has shown that macrophages appear to be required for the maintenance of the intestinal stem cell niche, where the ablation of intestinal macrophages resulted in decreased Lgr5 mRNA expression *in vivo* (Sehgal et al., 2018). Here, we show that the M1 macrophage population significantly upregulate colonic Lgr5+ stem cell numbers in a juxtracrine-contact dependent manner. In the small intestine, Lgr5+ stem cell expression is commonly regulated via neighboring Paneth cells providing essential factors such as Wnt3a, EGF, TGF-α and Notch ligand Dll4, however the colonic epithelium lacks Paneth cell expression and likely relies on external stimuli and subepithelial cues for maintenance of the stem cell niche (Sato et al., 2011; McCarthy et al., 2020). It is therefore possible that a cell-cell contact through Notch or the short-range Wnt signaling pathway may be involved in inducing the changes within the stem cell population we have observed (Farin et al., 2016; Sancho et al., 2015). Interestingly, earlier studies in human colonic epithelium have shown that Notch 1 was highly expressed in murine intestinal stem cells, while transcriptional profiling of bone-marrow derived M1 and M2 macrophages have demonstrated a significant increase in mRNA expression of the Notch ligands Delta-like ligand 1 and Jagged 1 in M1 macrophages when compared to non-activated (M0) and M2 macrophages (Li et al., 2018). The macrophages’ capacity to engage with the intestinal epithelium has been reported by numerous studies, especially in the colon, where most recently macrophages in the distal colon were shown to engage the epithelium through the formation of balloon-like protrusion used to limit the absorption of toxic fungal metabolites (Skoczek et al., 2014; Pull et al., 2005; Chakrabarti et al., 2018). As many of these findings are made in the large intestine, it is likely that the colonic epithelium is more receptive to macrophage-epithelial interactions compared to the small intestine, where the localization of Notch signaling receptors and ligands, specifically Jag-1, Dll-1, Dll-4 in the colon also differs to that of the small intestine (Shimizu et al., 2014; Noah and Shroyer., 2012; Van Dussen et al., 2012). The differential distribution of Notch ligands may allow macrophages with high Notch-signal receptor expression to engage and influence the intestinal stem cell niche within the colonic epithelium. However, to verify whether such a reciprocal interaction occurs *in vivo* further investigation is required (Ortiz-Masia et al., 2016; Monsalve et al.,2006).

Recent work in the small intestine has suggested that Paneth cell-derived Wnt3a is directly transferred to Lgr5+ stem cells therewith regulating the intestinal stem cell niche (Farin et al., 2016). As our findings indicate that the increase in Lgr5+ stem cell expression is dependent on M1-macrophage contact, it is feasible that a similar mechanism is utilized. In support of this hypothesis, we have demonstrated that juxtracrine interactions between M1 macrophages and the epithelium results in an upregulation of LEF1 and Cyclin D1, both of which are key downstream canonical Wnt signaling targets. Wnt signaling is often aberrantly dysregulated in chronic inflammatory bowel diseases where M1-like macrophages are abundantly localized (Hine et al., 2019). Our findings may add to the notion that the M1 macrophage phenotype likely contribute to increased Wnt signaling activity, thereby exacerbating intestinal disease progression (Moparthi and Koch., 2019; Cosin-Roger et al., 2019). Future work should aim to untangle the complex upstream Wnt signaling cascades involved which led in the increased activation of LEF1 and Cyclin D1 observed in our reductionist M1-crypt co-culture model.

Our study offers a first insight into the overarching effects of macrophage subtypes on colonic crypt proliferation and differentiation. Fundamental differences between the effects of M1 and M2 macrophages are particularly apparent on crypt stem cell driven differentiation. Strikingly, the physical contact of macrophages with the crypt epithelium can have profound effects on renewal that are distinct for each of the macrophage subtypes in this study. Here we have highlighted the significance of direct macrophage-epithelial contact and future studies should endeavor to delineate the complex signaling mechanism involved in the dialogue between the two parties, where this will provide fertile ground for the development of therapeutic strategies based on intestinal homing interference.

## Methodology

### *In Vitro* Experiments

All animal experiments were conducted in accordance with the Home Office Animals (Scientific procedures) Act of 1986, with approval of the University of East Anglia Ethical Review Committee, Norwich, U.K. Female C57BL/6 (UEA-Disease Modelling Unit) aged between 8-12 weeks, were euthanized by CO2 asphyxiation and subsequent cervical dislocation in accordance with Schedule 1 of the Act.

### Isolation and culture of bone marrow-derived macrophages

Following the isolation of the femur/tibia and the removal of residual connective tissue, the bone’s epiphyses were severed, and the bone marrow was flushed in a sterile environment using a 28-gauge syringe and cold RPMI 1640 (+10% FBS, +1% Pen/Strep). The flushed bone marrow contents were then then filtered through a 70μm nylon cell strainer (Falcon) and collected in a 50ml Centrifuge tube (Falcon). Following centrifugation at 600g for 10 minutes the cell suspension was re-suspended in warm RPMI1640. A bone-marrow yield was established, and the cells were appropriately seeded onto 6-well ultra-low attachment plates (Corning) at a cell density of 1 × 106! cells/ml. To drive BMDM differentiation towards macrophages, supplementary murine RM-CSF (Peprotech) at a concentration of 0.2μg/ml was added on Day 0 and 3 and macrophages were harvested on Day 8.

### Polarization of macrophage population

Macrophages were polarized based on methods previously described by Wei Ying et al in 2013 (Ying et al., 2013). BMDM cells were cultured in RPMI 1640 media up to day 7. On day 7, the floating cell population was removed, and the media was replaced by new fresh media. For M1 activation, supplementary LPS (100ng/ml) and IFN-γ (50ng/ml) were added to the media for a further 24 hours and for M2 activation, IL-4 (10ng/ml) and IL-13 (10ng/ml) were added instead.

### Isolation and culture of murine colonic crypts

Colonic crypts were isolated from the distal colon of C57BL6 mice, as previously described by Skoczek and colleagues (Skoczek et al., 2014). Briefly, following the culling of the mouse, the colon was removed and washed with ice-cold PBS to remove excess faecal matter; the colon was then cut longitudinally and excess mucus within the tissue was gently dissociated. 0.5mm tissue pieces were placed in a saline solution [50ml dH2O with NaCl (140mM), KCl (5mM), HEPES (10mM),d-glucose (5.5mM), Na2HPO4 (1mM), MgCl (0.5mM), CaCl (1mM), EDTA (1mM), DTT (0.153μg/ml), L-glutamine (200mM), Pen/Strep (200U/ml) and NEAA (2%)] for 1 hour. To liberate the crypts, the solution containing the tissue was shaken to aid gentle dissociation and then collected following crypt sedimentation. The single crypts were embedded in growth factor-reduced matrix Matrigel (VWR) and seeded onto No.0 glass coverslips (0.08-0.13mm) contained within 12-well plates (Starlab). Following polymerization of the Matrigel after 8 minutes at 37°C, the coverslips were flooded with colonic crypt culture media (Advanced DMEM/F12, containing B27 (20μl/ml), N2 (10μl/ml), N-acetyl-L-cysteine (0.163μg/ml).HEPES (10mM), Pencillin/Streptomycin (100U/ml), GlutaMAX (2mM), EGF (50ng/ml), Noggin (100ng/ml) (all from Peprotech), Wnt-3A (200ng/ml) and R-spondin-1 (1mg/ml) (BioTechne).

### Co-culture of macrophages and colonic crypts

To isolate the macrophages population, the cells were harvested on Day 8. On Day 8, the adherent population was liberated using 0.48 mM Versene. The optimum macrophage seeding density was previously determined to be 5.7×10^5^ cells per well which was then added to the colonic crypt/Matrigel solution. The mixture was then seeded onto a No.0 glass coverslip. Following Matrigel polymerization at 37oC, the Matrigel was then flooded with colonic crypt culture medium (as described above).

### Culture of colonic crypts with macrophage-conditioned media

Macrophages and crypts were isolated and cultured as previously described above. To study macrophage secretory factor-derived effects on colonic crypts, a three conditioned-media culture models were devised. Under the ‘control crypt’ model, two separated Matrigel’s with colonic crypts alone are seeded onto a well. Under the ‘M1 co-culture’ model, two separated Matrigel’s seeded with M1 macrophages and colonic crypts were seeded onto a well. Under the ‘M1 conditioned media (CM)’ model, two separated Matrigel’s with M1 macrophages seeded along with colonic crypts and another seeded with crypts alone was cultured. Under the ‘M1-only’ model, two separated Matrigel’s with colonic crypts seeded alone and another with M1 macrophage seeded alone is cultured onto a well. For EdU incorporation experiments the ‘control crypt’, ‘M1/M2-crypt co-culture’ and ‘M1/M2 conditioned media’ was utilized. The diagram in **Supplementary Fig 3** summarizes the experimental setup described above.

### Immunofluorescent labelling

For characterizing cells within the co-culture system, epithelial-specific antibodies were used. Following the co-culture, the coverslips were fixed with 4% PFA for 1 hour on ice. Washing steps were carried following each step. Ammonium chloride (100mM in PBS (pH7.4) was added to each coverslip for 13 minutes, washed with PBS, followed by further incubation with 10% SDS in PBS (pH 7.4) for 5 minutes. 1% Triton-X was added for 30 minutes to permeabilize the organoids. Non-specific binding was inhibited using 10% Donkey or Goat serum (Gibco) (depending on antigen retrieval) for 20 minutes.

Primary antibodies for enteroendocrine cells (Cro-A+) (Abcam), tuft cells (DCAMKL1+) (Abcam), Caspase 3 (Cell signaling) or stem cells (Lgr5+) (Origene) were added for overnight incubation at 4°C. The following day, immunolabelling was visualized using a species-specific Alexa-Fluor-conjugated secondary antibodies (488, 568, 647) raised in mouse, donkey, goat or rabbit and added for 2 hours at 4°C. PE-conjugated Ulex europaeus lectin (UEA-1) was acquired from VectorLabs to label goblet cells. Finally, the slides were washed and mounted with Hoechst/Vectashield (VectorLabs) the slides were later visualized using an epifluorescence or confocal microscopy.

### EdU incorporation experiments

Colonic crypts were cultured as previously described. Following 24 hours post-culture, EdU (10µM) was added to a final concentration of the crypts were then left to incubate at 37oC/5% CO2 overnight. On day 2, the crypts were fixed and processed as described above and EdU incorporation detected through a Click-iT reaction as per manufactures instructions (Thermofisher).

### Image Analysis

All fluorescent images were captured on the equatorial plane of the crypt as previously described (Jeffrey et al., 2016; Skoczek et al., 2014; Reynolds et al., 2013) using either a Nikon TI with a x20 0.4 NA, Zeiss Axiovert 200 with a x20 NA or Zeiss LSM-510-META with a x63 1.4NA 0.75mm WD oil immersion objective was used.

All images were analyzed with Fiji (Image J) software. To identify enteroendocrine cells (Cro-A+) (Abcam), tuft cells (DCAMKL1+) (Abcam), goblet cells (UEA-1+) (Vectorlabs), Caspase 3 (Cell signaling) and stem cells (Lgr5+) (Origene), Z-stacks were taken at a 1μm intervals for 5μm above and below the crypt equatorial plane. As luminal content is likely to appear as Caspase 3+, quantification of this marker was restricted to expression in cells within the crypt. Goblet cells were identified by following the UEA-1 + present within the cytoplasm of the individual cells throughout the equatorial plane. The location of each marker positive cell was recorded and separated into three crypt regions, base (cells within the +4 position of the crypt), mid and top region. To identify the stem cells within a crypt, the basal Lgr5 expression of each cell across the Z-stack (optical slices), 5μm above and below, the equatorial plane was counted. Crypt budding numbers were quantified by counting the buds present on day 1 and day 6 of culture.

### Quantification of nuclear fluorescence intensity

Images were captured using the confocal microscope (LSM-510-META) with a x63oil a x63 1.4NA 0.75mm WD oil immersion objective. To quantify the expression of Cyclin-D1 and LEF1 within the nucleus, the average fluorescence value of every nucleus present at the equatorial plane was measured. Using Fiji-Image J’s polygon tool, the nuclear area was identified by following the perimeter of each individual DAPI+ nuclei in the equatorial plane. The arbitrary fluorescent value of the channels occupied by Cyclin-D1 and LEF1 were then measured as shown in **Supplementary Fig 4**.

### Statistical Analysis

All experiments were repeated at least three times unless stated otherwise. Data are expressed as mean ± standard error of mean (SEM), n= number of independent experiments, N= minimum total number of crypts and a minimum of 20 crypts per experiment were counted. Statistical analysis was carried out using the Graphpad Prism 9 software. Comparisons between two or more groups were measured using one-way ANOVA with post-hoc Tukey analysis and a paired t-test was utilized to compare differences between two groups. P-value of less than 0.05 was considered statistically significant.

