## Supplementary Materials for "M1 and M2 macrophages differentially regulate colonic crypt renewal"

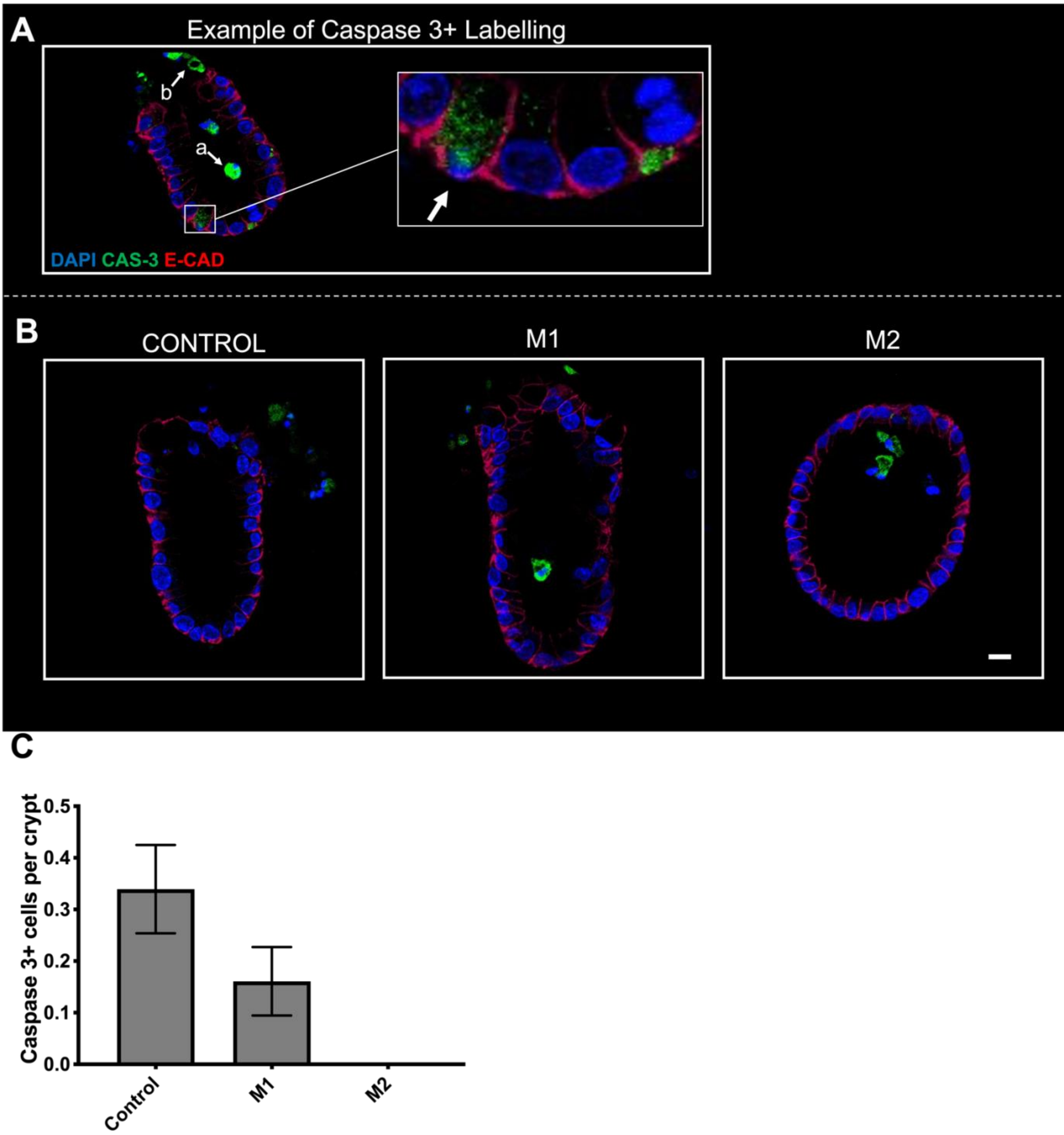


***Supplementary Figure 1: Expression of Caspase-3 in the colonic epithelium within a 3D macrophage-crypt co- culture.***

***A)*** *An example of a Caspase-3 positive cell (white arrow) expressed within the crypt epithelium. Caspase-3 expression present in the lumen and shedding region in the upper region of the crypt but are distinctly absent in cells within the crypt.*

*Positive example shown above (lumen- **a** and shedding area-**b**). **B)** Representative confocal images showing Caspase-3 expression (green), nuclei (blue) and E- cadherin (red) in crypt-macrophage subtype co-culture. Scale bar at 15μm. **C)** Histogram showing the average number of Caspase-3 positive epithelial cells per crypt within each co-culture condition (n=3, ns).*

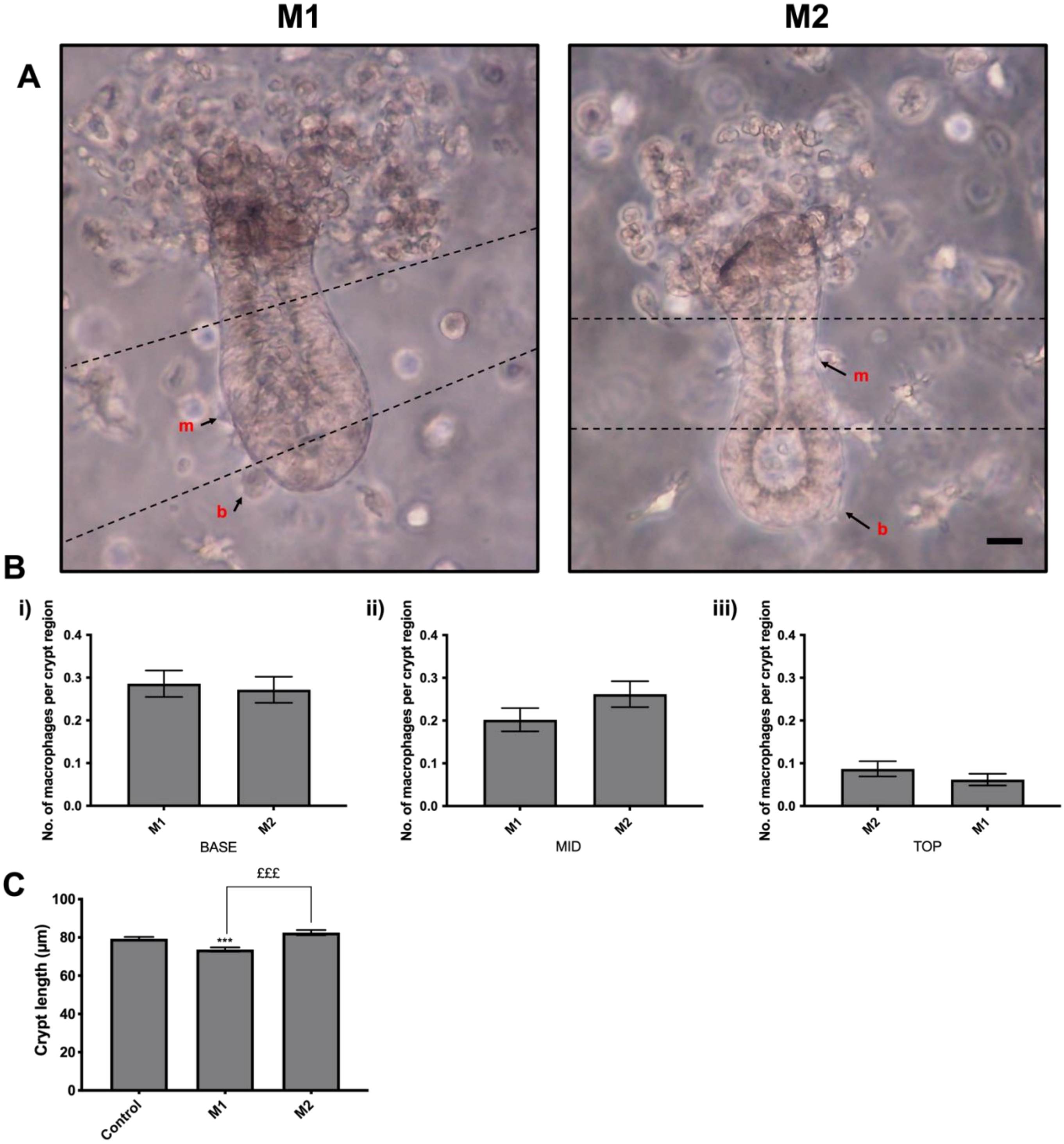

***Supplementary Figure 2: Macrophage-crypt contact is maintained in the presence of either M1 or M2 macrophages in vitro*** ***A)*** *Representative brightfield images showing macrophages in contact at the base (**b**), mid (**m**) regions of the crypt after 24 hours in vitro culture* ***B)*** *Histogram showing the average number of macrophages distributed along each crypt region* ***i)*** *base,* ***ii)*** *mid and* ***iii)*** *top (n=10; *P<0.05 compared to Control).* ***C)*** *Histogram showing the average length of crypts in culture with M1 or M2 macrophages after 24 hours (n=6; $^{***}$P<0.001 compared to Control; $^{£££}$P<0.001 M1 compared to M2). Scale bar at 15µm.*

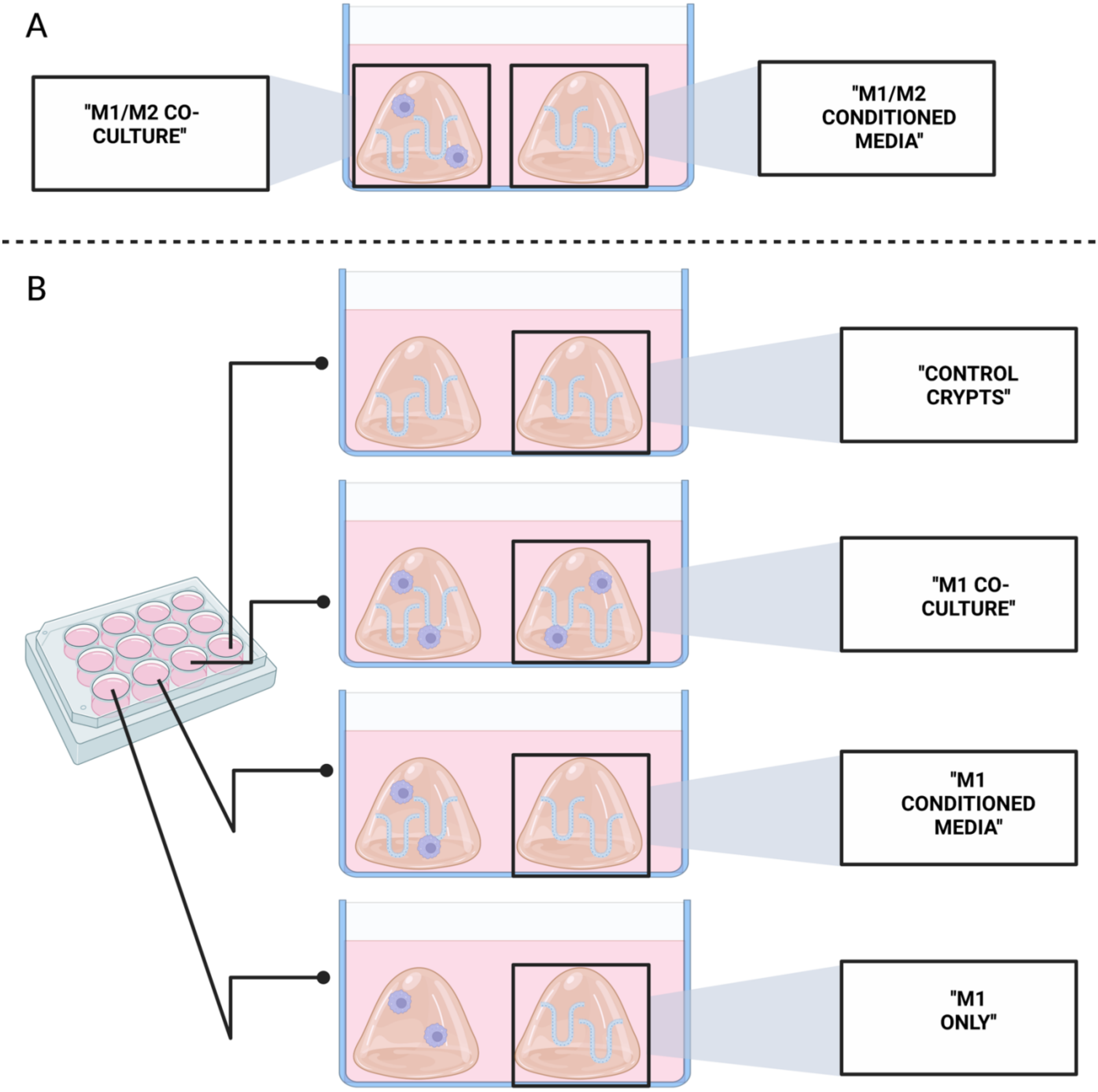

***Supplementary Figure 3: Conditioned media experimental setup***

*Diagram showing the experimental setup for the study of physical vs secretory factors* ***A)*** *M1 and M2 macrophage conditioned media model utilized for EdU incorporation experiments* ***B)*** *Model of M1 physical vs secretory experiment. Under the 'control crypt' model, two separated Matrigels with colonic crypts alone are seeded onto a well. Under the 'M1 co-culture' model, two separated Matrigel's seeded with M1 macrophages and colonic crypts were seeded onto a well. Under the 'M1-CM (conditioned media)' model, two separated Matrigel's with M1 macrophages seeded along with colonic crypts and another seeded with crypts alone was cultured. Under the 'M1-only' model, two separated Matrigel's with colonic crypts seeded alone and another with M1 macrophage seeded alone is cultured onto a well. Created in Biorender (2022).*

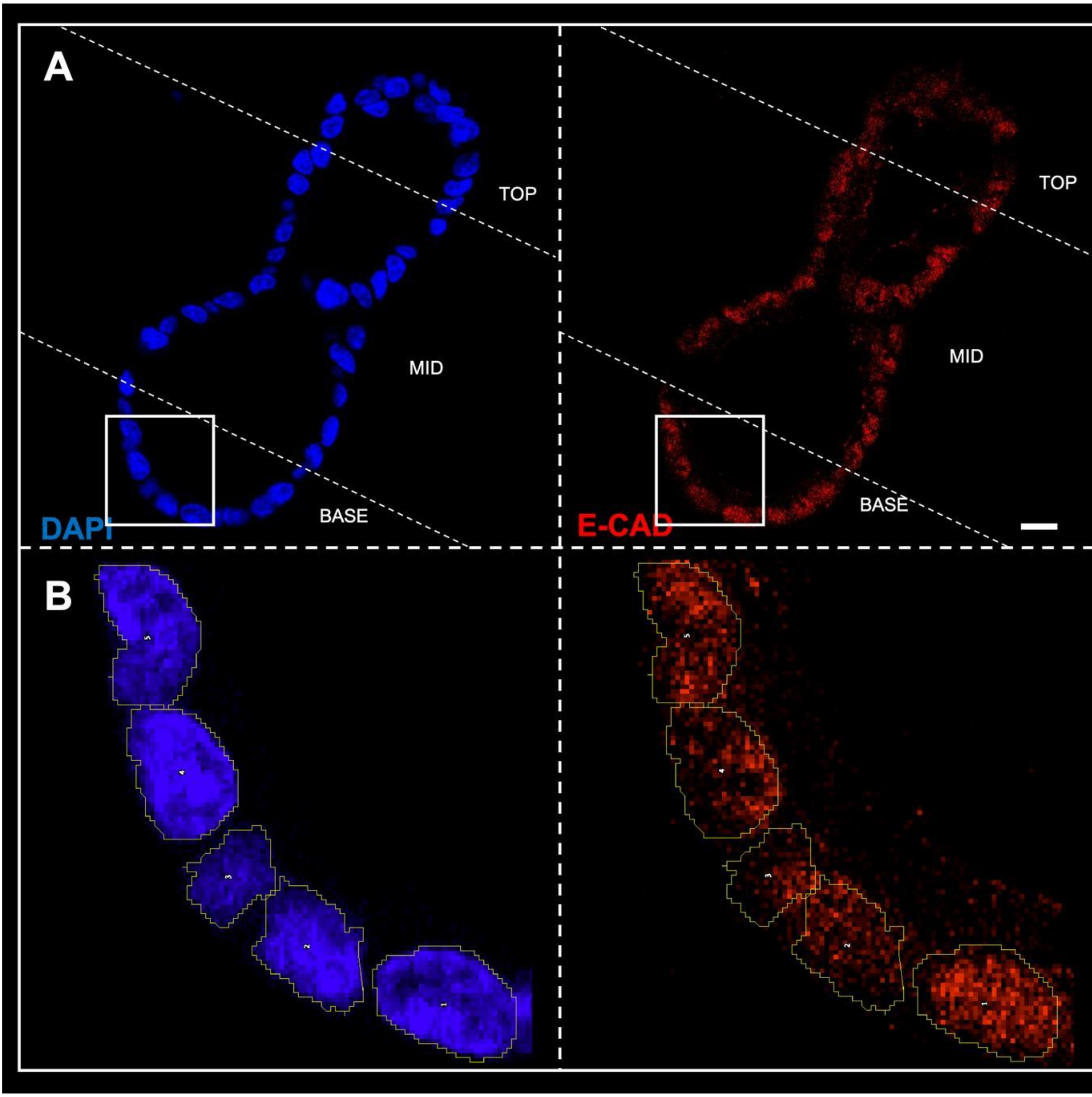


***Supplementary Figure 4: Identification and analysis of nuclear LEF1 localization***

*Analysis of nuclear fluorescence intensity per crypt* ***A)*** *DAPI+ and LEF1 expression in a crypt* ***B)*** *The nuclear area was highlighted using DAPI and the channel was switched to LEF1, where the average fluorescence intensity of the area was measured. This analytical approach was also mirrored with analysis of Cyclin D1. Scale bar at 15µm.*
